## Supplemental Figures for "Cas9 targeted enrichment of mobile elements using nanopore sequencing"

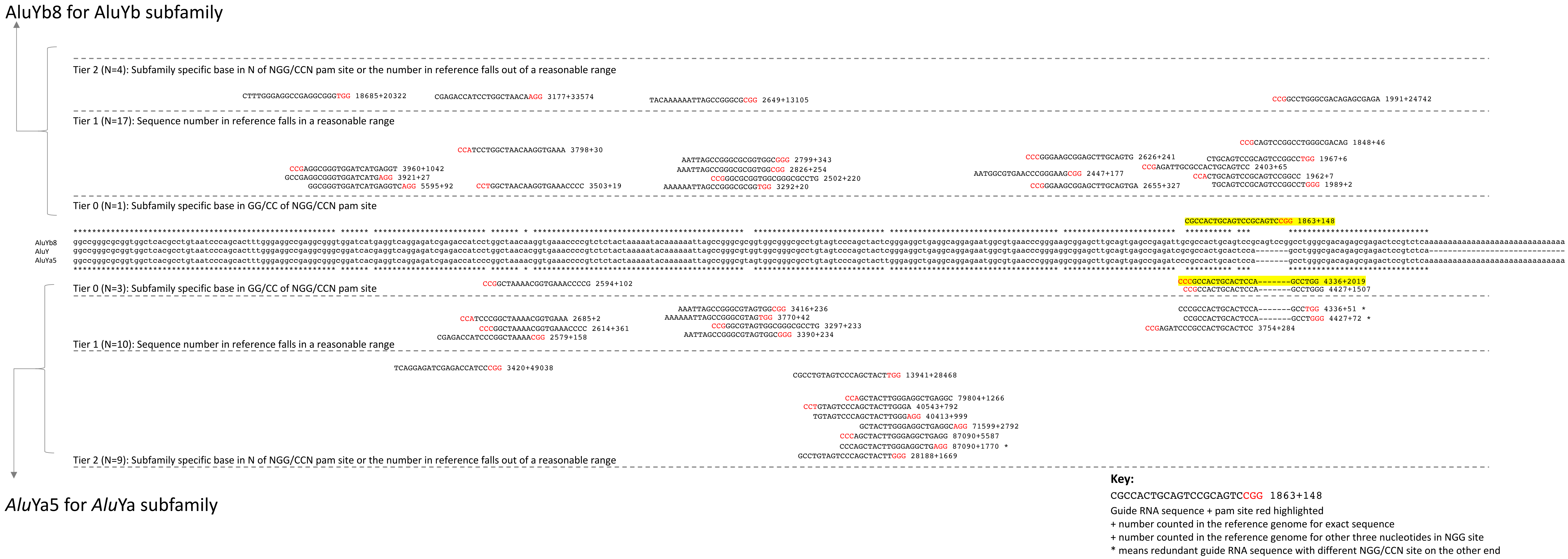


**Supplementary Fig.1: Guide RNA design for *Alu*Yb and *Alu*Ya element.**
Consensus sequences are aligned and showed for *Alu*Yb, *Alu*Ya, and *Alu*Y in the middle. The candidate guide RNAs are distributed based on different tiers (see **Methods**). The final list is highlighted by yellow.

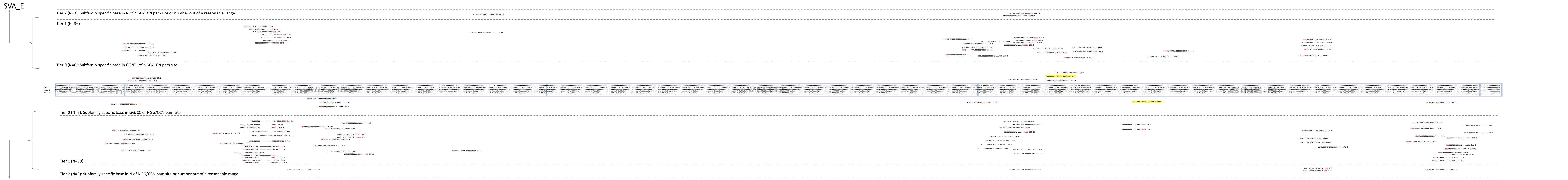


**Key:**

CGCCACTGCAGTCCGCAGTCCGG 1863+148

Guide RNA sequence + pam site red highlighted

+ number counted in the reference genome for exact sequence

+ number counted in the reference genome for other three nucleotides in NGG site

* means redundant guide RNA sequence with different NGG/CCN site on the other end

**Supplementary Fig.2: Guide RNA design for SVA_F and SVA_E element.**
Consensus sequences are aligned and showed for SVA_F, SVA_E, and SVA_D in the middle. The candidate guide RNAs are distributed based on different tiers (see **Methods**). The final list is highlighted by yellow.

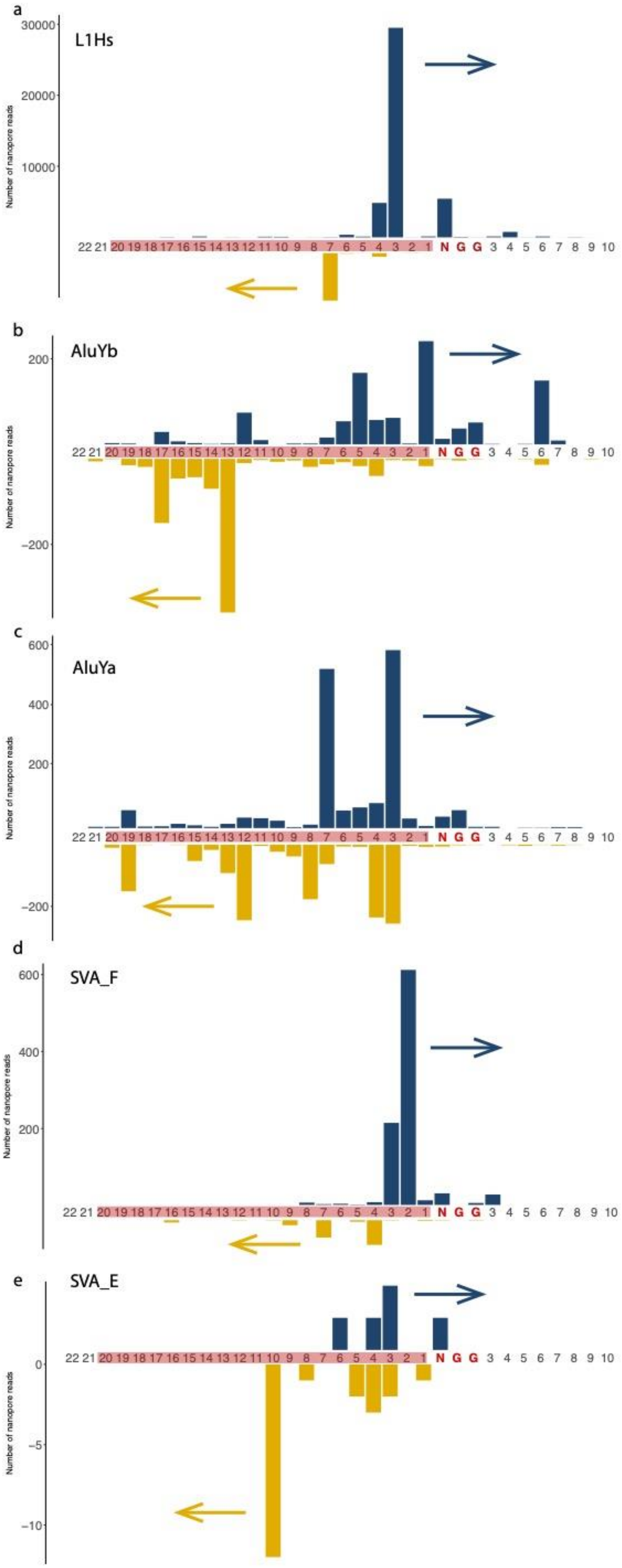
a
L1Hs
30000
20000
10000
Number of nanopore reads
22 21 20 19 18 17 16 15 14 13 12 11 10 9 8 7 6 5 4 3 2 1 N G G 3 4 5 6 7 8 9 10
b
AluYb
200
−200
Number of nanopore reads
22 21 20 19 18 17 16 15 14 13 12 11 10 9 8 7 6 5 4 3 2 1 N G G 3 4 5 6 7 8 9 10
c
AluYa
600
400
200
−200
Number of nanopore reads
22 21 20 19 18 17 16 15 14 13 12 11 10 9 8 7 6 5 4 3 2 1 N G G 3 4 5 6 7 8 9 10
d
SVA_F
600
400
200
Number of nanopore reads
22 21 20 19 18 17 16 15 14 13 12 11 10 9 8 7 6 5 4 3 2 1 N G G 3 4 5 6 7 8 9 10
e
SVA_E
0
−5
−10
Number of nanopore reads
22 21 20 19 18 17 16 15 14 13 12 11 10 9 8 7 6 5 4 3 2 1 N G G 3 4 5 6 7 8 9 10

**Supplementary Fig.3: Distributions of guide RNA cleavage-site for five MEI subfamilies**

**a,** Cleavage-site distribution of L1Hs guide RNA. X-axis shows the position where the read ends or begins with the number indicating the distance from the 'N' of the PAM site (NGG). The PAM site (NGG) was colored red and guide RNA bases were highlighted by red background. Y-axis is the number of nanopore reads counted. The upper blue bar represents the reads with forward strand sequencing outward the 3' end of guide RNA and the lower maize bar represents the reads with reverse strand sequencing outward the 5' end of guide RNA. **b,** Cleavage-site distribution of *Alu*Yb guide RNA. **c,** Cleavage-site distribution of *Alu*Ya guide RNA. **d,** Cleavage-site distribution of SVA_F guide RNA. **e,** Cut-site distribution of SVA_E guide RNA.

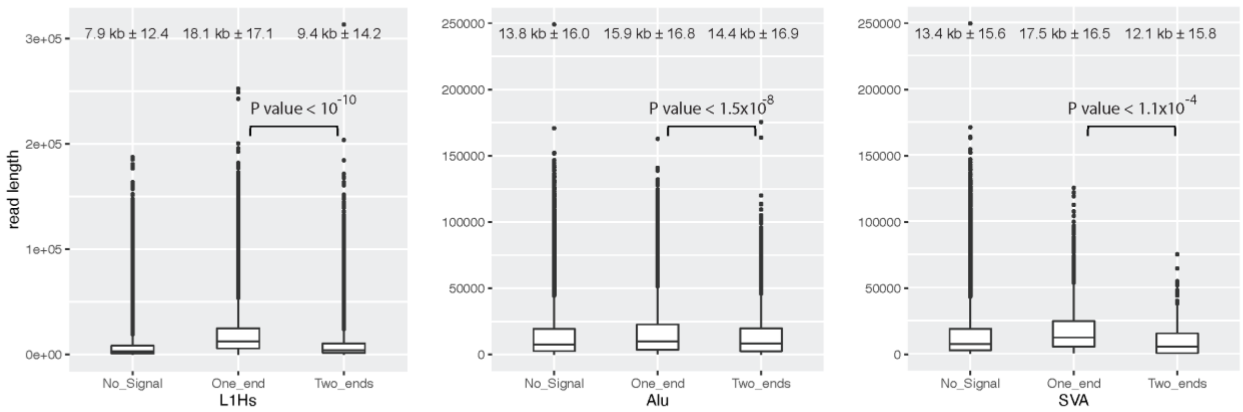


**Supplementary Fig.4: Read length distributions for MEI categories (L1Hs, *Alu*Y, and SVA).**

All reads were identified into three categories: reads with on-target reference MEI signals, ones with on-target non-reference MEI signals, and ones with no signals (off-targeted). The read length (mean ± standard deviation) are shown above the boxplots. And the P-value (student's T-test, two-tailed) are shown between the boxplots of one-end reads and two-end reads. Error bars range from Q1-1.5IQR to Q3+1.5IQR (IQR, interquartile range).

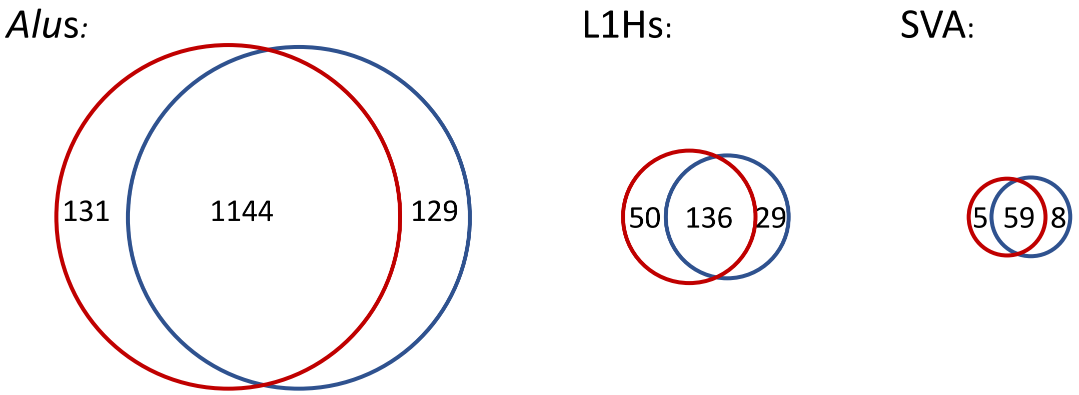


**Supplementary Fig.5: Venn diagram of the PALMER callset and the PAV callset for non-reference MEIs in NA12878 genomes.**

PALMER callset (red circle) is from PacBio raw sub-reads, and PAV callset (blue circle) is from PacBio assembly-based pipeline. The circles are depicted by the scale of the numbers showed inside. The union of two sets are generated as ‘PacBio-MEI’ to be the gold standard set to compare with the calls from nanopore data.

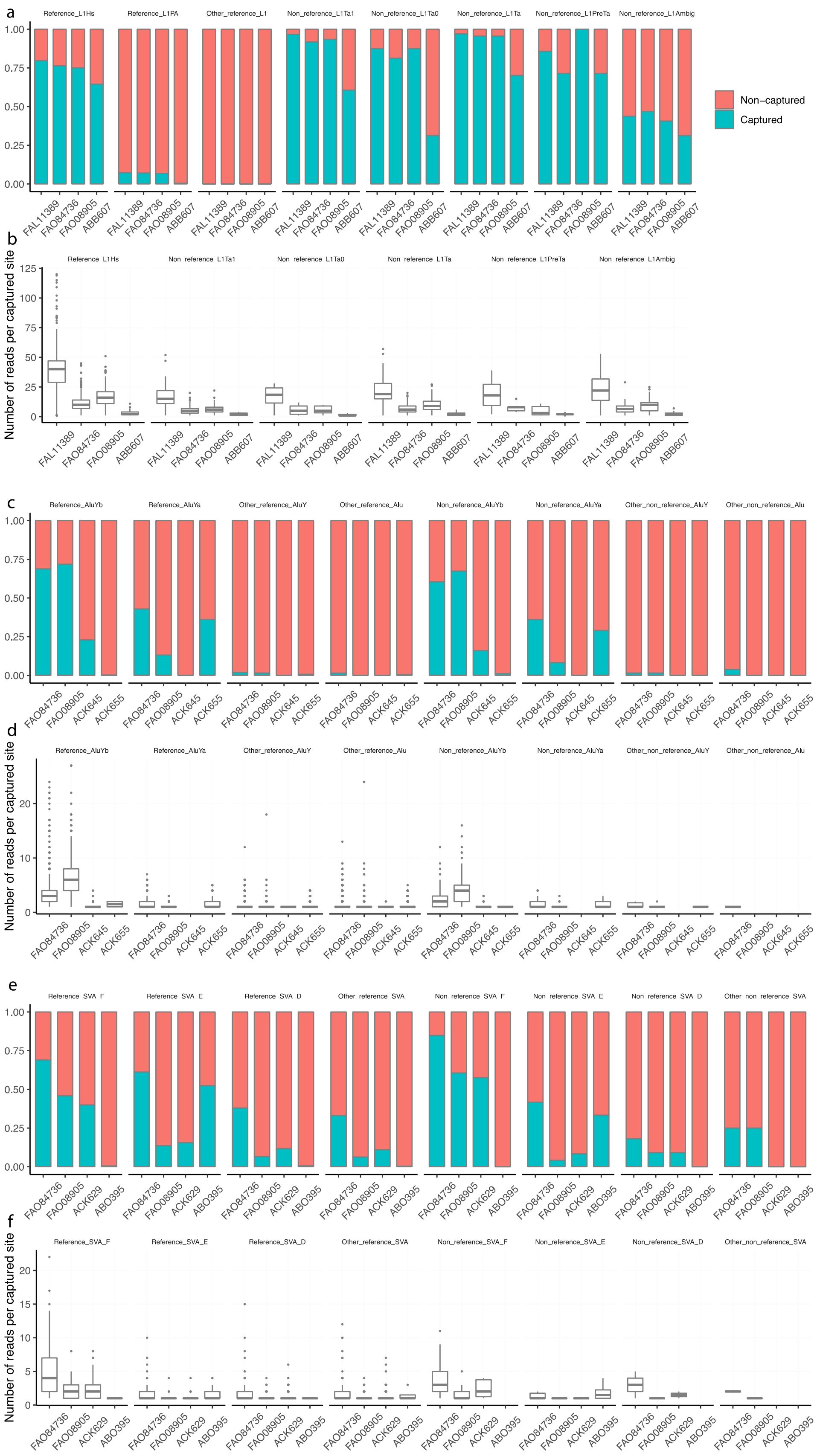
a
Reference_L1Hs
Reference_L1PA
Other_reference_L1
Non_reference_L1Ta1
Non_reference_L1Ta0
Non_reference_L1Ta
Non_reference_L1PreTa
Non_reference_L1Ambig
Non−captured
Captured
FAL11389
FAO84736
FAO08905
ABB607
b
Number of reads per captured site
c
Reference_AluYb
Reference_AluYa
Other_reference_AluY
Other_reference_Alu
Non_reference_AluYb
Non_reference_AluYa
Other_non_reference_AluY
Other_non_reference_Alu
ACK645
ACK655
d
e
Reference_SVA_F
Reference_SVA_E
Reference_SVA_D
Other_reference_SVA
Non_reference_SVA_F
Non_reference_SVA_E
Non_reference_SVA_D
Other_non_reference_SVA
ACK629
ABO395
f

**Supplementary Fig.6: Summary of recovered known reference and non-reference MEIs.**

**a,** Known L1Hs in GM12878 recovered by Cas9 targeted enrichment from the individual MinION flow cell (FAL11389), pooled-MEI MinION flow cell (FAO84736), and individual Flongle flow cell (ABB607), displayed in a way of proportion of the upper-bound known reference L1Hs, L1Pa, and other L1 as well as non-reference (non-ref.) subfamilies (L1Ta1, L1Ta0, L1Ta, L1PreTa, and L1Hs with ambiguous subfamilies) of L1Hs from PacBio-MEI set. **b,** The number of supporting reads in each captured L1 in the context of **a.** **c,** Known *Alu*Y elements in GM12878 recovered by Cas9 enrichment in two pooled MinION flow cells (FAO84736 and FAO08905), one individual *Alu*Yb Flongle flow cell (ACK645), and one individual *Alu*Ya Flongle flow cell (ACK655). **d,** The number of supporting reads in each captured *Alu* element in the context of **c**. **e,** Known SVA elements in GM12878 recovered by Cas9 enrichment in two pooled MinION flow cells (FAO84736 and FAO08905), one individual SVA_F Flongle flow cell (ACK629), and one individual SVA_E Flongle flow cell (ACK395). **f,** The number of supporting reads in each captured *Alu* element in the context of **e**. Error bars of boxplot range from Q1-1.5IQR to Q3+1.5IQR (IQR, interquartile range).

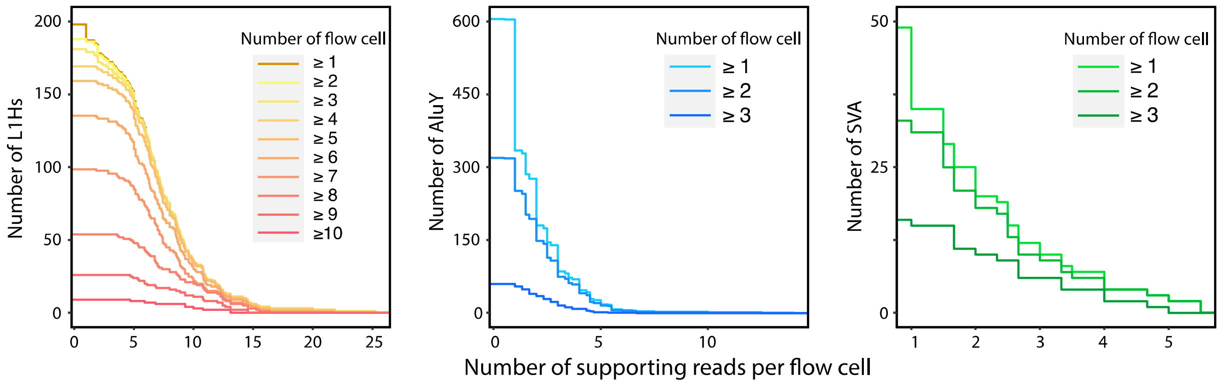


**Supplementary Fig.7: MEI distributions in various number of flow cells.**

The number of MEIs (L1Hs, yellow; *Alu*Y, blue; SVA, green) can be captured by nanopore Cas9 enrichment regarding different numbers of flow cells and cutoffs of supporting reads.

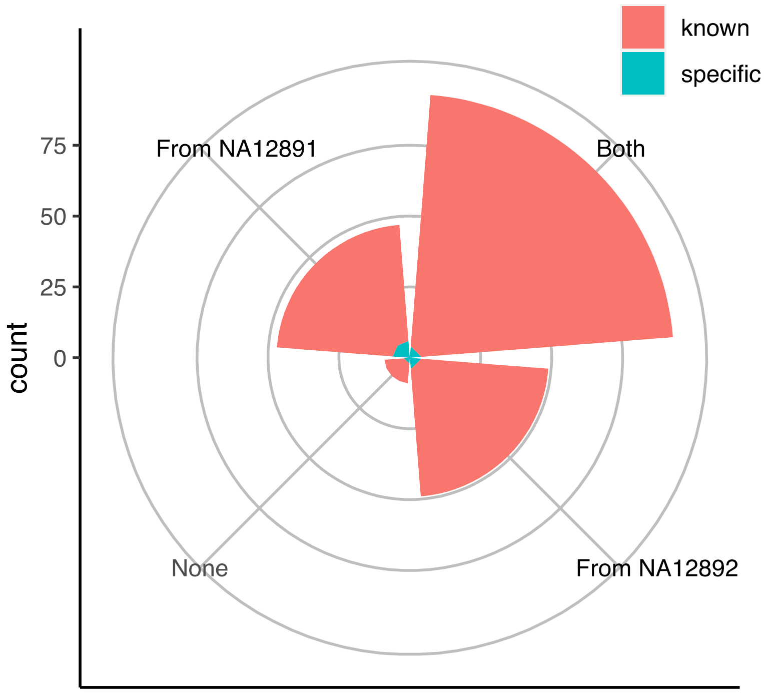


**Supplementary Fig.8: Trio transmission of 198 non-reference L1Hs captured by nanopore in GM12878 sample.**

The intersections with ‘PacBio-MEI’ were shown by red and the nanopore-specific non-reference L1Hs that were missed by ‘PacBio-MEI’ were shown by green.

**Supplementary Table 1. A list of guide RNA candidates**

Sheet1, five final guide RNAs in the project for five MEIs, along with the information of the counts of forward/reverse strand reads based on these guide RNA sequence. Sheet2, all candidate guide RNAs for L1Hs that mapping to unique sequences of L1Hs subfamily followed by the frequency in the reference genome sequence. Sheet3 for *Alu*Yb, Sheet4 for *Alu*Ya, Sheet5 for SVA_F, and Sheet6 for SVA_E.

**Supplementary Table 2. Counts of forward and reverse reads in cleavage-site analysis.**

**Supplementary Table 3. Master table of Information for all flow cells in the project**

Sheet1, information of reads from all 17 flow cells. Sheet2, information of reads after MEI classification by Nano-Pal. Sheet3, number of MEI events captured by each flow cell after reads clustering.

**Supplementary Table 4. Non-reference MEI callsets from PacBio-MEI and MELT.**

Sheet1, PacBio-MEI L1Hs callset in GM12878. Sheet2, PacBio-MEI *Alu* callset in GM12878. Sheet3, PacBio-MEI SVA callset in GM12878. Sheet4, MELT L1Hs callset for GM12891. Sheet5, MELT L1Hs callset for GM12892.

**Supplementary Table 5. Known MEIs captured by nanopore Cas9 enrichment approach in different flow cells based on different boundaries.**
Upper-bound, intermediate, and lower-bound values of different categories of MEIs are included regarding background (number) and seven representative flow cells(percentage).

**Supplementary Table 6. Enrichment of mobile element signals in nanopore reads from GM12878 trio L1Hs experiments**
Four flow cells were carried out for trio experiments in the project: three individual Flongle flow cells for GM12878 (ABG188), GM12891 (ABO515), and GM12892 (ABN780) each, and one MinION flow cell for pooled three samples (FAL15177).

**Supplementary Table 7. Information for nanopore-specific captured non-reference MEIs and examples**

Sheet1, summary of potential nanopore-specific non-reference MEIs in different categories (see **Methods**). Sheet2, information of potential nanopore-specific non-reference L1Hs. Sheet3, information of potential nanopore-specific non-reference *Alu*Y elements. Sheet4, information of potential nanopore-specific non-reference SVAs. Sheet5, IGV screenshots for an L1Hs example at chrX:121709062-121709136 as true positive. Sheet6-8, IGV screenshots of examples of L1Hs identified as false positives. Sheet9,10, IGV screenshots for examples of *Alu*Ys identified as true positives. Sheet11-13, IGV screenshots for examples of *Alu*Ys and SVAs identified as false positives.

**Supplementary Table 8. MEI callset by Cas9 targeted enrichment using nanopore sequencing in GM12878**

Sheet1, information for the L1Hs callset in GM12878. Sheet2, information for the *Alu*Y callset in GM12878. Sheet3, information for the SVA callset in GM12878.
